# Effects of evolution on niche displacement and emergent population properties, a discussion on optimality

**DOI:** 10.1101/2020.05.04.075994

**Authors:** Rudolf P. Rohr, Nicolas Loeuille

## Abstract

Understanding the effects of evolution on emergent population properties such as intrinsic growth rate, species abundance, or dynamical resilience is not only a key theoretical question, but has major empirical implications for conservation, agroecology, invasion ecology among others. In particular, could we classify evolutionary scenarios leading to optimisation of those properties, from the ones who do not. First, we uncover two classes of invasion fitness functions, only the first one allowing optimization of some (but typically not all) population properties. Second, we showed that our two classes are also strongly linked to niche displacement and emergence of polymorphism. Our results indicate that optimization is, in general, incompatible with niche differentiation and, therefore, with emergence of polymorphism through evolutionary branching. Actually, niche displacement between resident and mutant morphs, and potentially polymorphism, only arise when we do not expect optimality to hold. We extensively discuss which biological traits can fall into which class of invasion fitness. Although, it is possible to find traits for which optimality is expected, we argue that for the majority of the cases it does not hold. Finally, we provide practical applications of our results in conservation, agroecology, harvesting and invasion ecology.

## Introduction

Recently, empirical results have accumulated, suggesting that evolution can often impact the dynamics of ecological systems, even on short timescales (Hairston et al., 2005). As a result, we may have to account for such evolutionary phenomena to properly predict or manage the effects of current global changes (Urban et al., 2016; Carroll et al., 2014), to manage exploited species (Allendorf et al., 2008) or agricultural systems (Loeuille et al., 2013; Hendry et al., 2011). Understanding the ecological consequences of evolution requires a non trivial change in scale, from genes and individuals to populations and ecosystems. For instance, natural selection builds on differences in fitness components of different genotypes. How such individual-level differences scale up to influence the fate of populations is already quite a difficult question. While evolution may enhance population persistence by fostering adaptation (e.g., evolutionary rescue: Gomulkiewicz and Holt (1995)), it may also lead to the fixation of traits that decrease population size or even lead to extinctions (Webb, 2003). Eco-evolutionary dynamics also alter ecological properties at even larger scales, affecting community structure (e.g., competitive hierarchies: De Meester et al. (2002)) and ecosystem functioning (e.g., the strength of trophic cascades and nutrient recycling processes: (Loeuille and Loreau, 2004; Bassar et al., 2010)).

Evolution has often been construed as positive for the evolving population. Statements on evolution happening “for the good of the species” have been common, giving the impression that evolution optimizes various aspects of population properties (e.g., its abundance or productivity). Many models in evolutionary biology support this view. This may come from a strong focus on adaptation compared to other evolutionary processes, and to the fact that frequency dependence is often disregarded. As explained in Dieckmann and Ferrière (2004), many quantitative genetic models are based on ad-hoc fitness functions that ignore frequency dependent selection and such models often lead to an evolutionary outcomes that optimize intrinsic growth rates or reproduction ratio *R*_0_.

Similar ideas of optimization have also been proposed at ecosystem scales, where some works have proposed that ecosystems naturally evolve toward optimal states (Odum and Barrett, 1971). In recent years, however, several works have shown that evolution only rarely optimizes systems at any of these scales (populations or ecosystems). Dieckmann and Ferrière (2004) discuss how frequency-dependence naturally emerges from the ecological context and will most often lead to evolutionary outcomes that do not optimize population characteristics. Metz et al. (2008) show that optimization through evolution is only obtained when the link between fitness and the environment is simple (unidimensional).

When and whether evolution optimizes ecological properties of natural populations also has important implications from an applied perspective. Let us consider the link between evolution and the growth rate of species. Understanding when evolution leads to maximal intrinsic growth rates has important implications for the conservation of rare species, whose growth rate restoration is a priority. Evolution also plays a key role in the propagation of many invasive species (Mooney and Cleland, 2001; Phillips et al., 2008, 2006; Shine et al., 2011). Because invasive species management partly aims at decreasing the invasive species population growth rate, understanding how observed evolutionary changes impact the species growth rate can have far reaching implications. Now consider the link between evolution and standing biomass or density. Understanding this link has important implications for the management of exploited species and agricultural systems. In fisheries for instance (Olsen et al., 2004; Conover and Munch, 2002), but also in hunts (Coltman et al., 2003), fast evolution results either from the strong mortality pressures exerted on the exploited populations or from targeting specific phenotypes (e.g., harvesting regulation based on fish size). How evolutionary changes alter final (equilibrium) biomass constrain our future management of exploited stocks. In fisheries, evolution toward earlier maturity or smaller body sizes has repeatedly been reported with large implications for sustainability (Conover, 2000). In agriculture, humans act as direct selective forces, affecting the evolution of wild species (e.g., evolution of pesticide resistance (Carlson et al., 2014)), and sorting cultivated species and phenotypes at different scales (artificial selection). Knowing which evolutionary scenarios enhance or optimize standing biomass would open new doors for a sustainable agricultural management (Denison et al., 2003; Loeuille et al., 2013). Finally, consider the link between evolution and system resilience. Previous works suggest that evolution likely alters ecosystem resilience (Loeuille, 2010; Kondoh, 2003), possibly reducing it in diverse communities (Loeuille, 2010). Eco-evolutionary dynamics also affect the occurrence of regime shifts in natural systems (Dakos et al., 2019). Again, understanding the conditions allowing very resilient systems will likely be important in a world that currently undergoes large disturbances (Carlson et al., 2014; Urban et al., 2016; Carroll et al., 2014).

To better understand the link between evolution and various ecological properties, one possibility is to understand the variations in the mean phenotype, and to explicitly account for impacts of such phenotypic variations on the ecological dynamics. However, it is also possible that evolution leads to character displacement within the population and to the long term maintenance of polymorphisms (Doebeli and Dieckmann, 2000; Leimar, 2005). Considering the mean phenotype is then no longer relevant. As character displacement occurs, due to disruptive selection, different phenotypes exploit different niches. This not only limits competition and related losses, but also leads to higher complementarity in resource use. These two components can act in synergy to allow higher abundances or productivity to be reached. Therefore, when niche displacement occurs, we expect some of the population characteristics to be positively affected, as competition is relaxed.

In this article, we start by investigating how evolution simultaneously changes three emergent population properties: the population intrinsic growth rate, its standing biomass and its resilience. Then, by integrating coexistence theory (Chesson, 1990; Saavedra et al., 2017) and eco-evolutionary dynamics, we study how character displacement and the emergence and maintenance of polymorphism affects these evolutionary consequences. To do this, we rely on a simple Lotka-Volterra competition model. We distinguish different evolutionary scenarios changing in the components of the model affected by the phenotype. More specifically, we have four aims. First, from a conceptual of view, we would like to uncover scenarios that lead to optimization of the different properties we consider. Second, we study whether optimization happens simultaneously for the three emergent properties, or whether evolution actually optimize a subset of these, at the expense of other emergent properties (e.g., leading to a very productive, high biomass system but with little resilience). Third, by integrating the concept of niche differentiation into eco-evolutionary dynamics, we aim at studding how niche/no-niche displacement relates to the optimization of the three emergent properties and to the emergence and enhancement of polymorphism. Finally, our aim is also to propose biological traits that would fit these different scenarios. This classification would help us, when strong selection is reported for a given biological trait, to propose how the resulting evolution may qualitatively alter various population characteristics that matter from a management point of view. We mathematically analyze these more complex scenarios, and illustrate coevolution scenarios using numerical simulations. Our analysis shows that evolution optimizes emergent properties for traits that solely impact fecundity or survival or traits that offer *per se* a competitive advantage. But, at the same time, there is no-niche differentiation between resident and mutant, consequently, no emergence and maintenance of polymorphism through branching. For traits whose effects on competition are context dependent (e.g., depend on traits of other individuals within the population), optimization is unlikely, and will not happen for all emergent properties. However, emergence of polymorphism through branching, due to niche differentiation, can occur. Because we find that many traits in fact fit this latter scenario, we propose that optimization will not often be observed.

## Main result

Within the adaptive dynamic framework, we define two main classes of relative fitness functions. These two classes have different consequences on the optimization of emergent population properties, on the type of evolutionary singular strategies, and on the niche differentiation between the resident and the mutant morphs. Adaptive dynamics considers explicitly the feedback loop between the ecological and the evolutionary dynamics by considering density and frequency dependent selection (Dieckmann and Law, 1996). It is therefore perfectly tailored to study to what extent evolution can optimize emergent population properties such as intrinsic growth rate, abundance, and dynamical resilience. In adaptive dynamics, the direction of evolution is determined by the so-called relative fitness function, which is based on the invasion fitness of a rare mutant of trait *x*_*m*_ in a resident population of trait *x* (Metz et al., 1992). *In turn, the invasion fitness is explicitly determined by the ecological dynamics and it is computed as the per capita* growth rate of a rare mutant in a resident population, considered at its ecological equilibrium *N*\*(*x*). The mutant *x*_*m*_ can invade if the invasion fitness is positive. Assuming small mutations, evolutionary changes are then proportional to the fitness gradient assessed at the resident phenotype, and evolutionary singularities correspond to the roots of the fitness gradient (Dieckmann and Law, 1996).

We study how evolutionary dynamics affect three emergent population properties. The first is the intrinsic growth rate or the Malthusian growth rate *r*, which is computed as the per capita growth rate when the population abundance tends to zero. The second is the abundance at equilibrium *N*\*. And the third is the dynamical resilience *λ* define as the return rate to equilibrium after small perturbation in abundance. In general it is computed by the real part of the leading eigenvalue of the Jacobian matrix, which in the case of a single population equals its intrinsic growth rate.

The relative fitness function can be classified into two classes defined by the following equations:

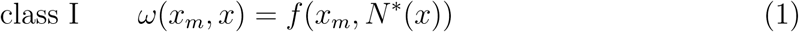

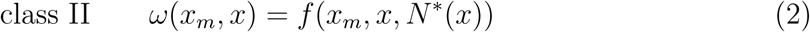

In the first class (equ. 1), the invasion fitness is a direct function of the mutant trait *x*_*m*_ only. The traits of the resident *x* acts only through the ecological equilibrium of the resident population *N*\*(*x*). While in the second class (equ. 2), the invasion fitness is a direct function of both the resident and the mutant traits. As we will show this subtlety between class I and class II leads to completely different results in the type of evolutionary singular strategies, niche/no-niche differentiation, and affects the link between evolution and the three studied emergent ecological properties.

Class I leads to singular strategies that can only be convergent and non-invasible (CSS) or non-convergent and invasible (repellor). Moreover, we show that CSS cases correspond to a local maximum in species abundance, while repellors correspond to a local minimum in species abundance, so that evolution always increases population abundances in this class of model. In class I, intrinsic growth rate or dynamical resilience may also be optimize, but this is not a general rule. In turn, class II can lead to any type of singular strategies, can create niche differentiation between the resident and the mutant morphs, and evolution does not optimize the three emergent population properties, in general.

We will start by illustrating our main results in a series of three studied cases, going from the optimization of all three emergent ecological properties, through the optimization of abundance only, and finally the case of non-optimization. Then, we will give an ecological interpretation of the two classes of relative fitness function in terms of niche/no-niche differentiation. We also discuss biological traits or situations that may fit one class or the other. It will be followed by the mathematical proof of our main results. Finally, we will generalize our results to the case of polymorphism (or species) coevolution.

### Optimization of all three emergent population properties

As a first study case, we consider ecological dynamics following the classical Verhulst model. Moreover, we assume that only the Malthusian intrinsic growth rate *r*(*x*) > 0 is function of the adaptive trait *x*. Population dynamics of trait *x* are then given by

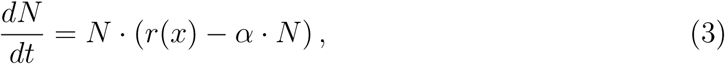

where the parameter *α* > 0 is the intraspecific competition. In this simple population model, the invasion fitness of a rare mutant *x*_*m*_ in a resident population at ecological equilibrium *N*\*(*x*) = *r*(*x*)/*α* is given by

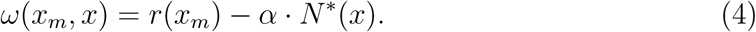

This function is clearly a relative fitness of class I and evolution optimizes species abundance. In this case, evolution also optimizes the Malthusian intrinsic growth rate *r*(*x*) and the dynamical resilience (which in this case equals the intrinsic growth rate *λ*(*x*) = *r*(*x*)). These various optimizations can be proved by studying the fitness gradient. In this case, it is directly proportional to the ecological equilibrium abundance gradient and to the intrinsic growth rate gradient:

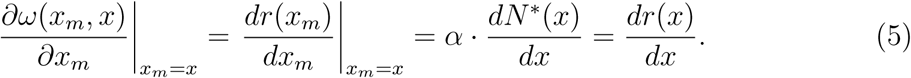

Therefore, singular strategies, i.e., roots of the fitness gradient, are also extrema in abundance and in intrinsic growth rate. Evolutionary singular strategies can be either convergent (in which case local trajectories will converge to the strategy) or divergent (in which case natural selection favors strategies away from the singular strategy). They can also be invasible by nearby mutants, or non-invasible (ESS). Convergence and invasibility of singular strategies can be investigated using second derivatives of the relative fitness, assessed at the singularity (Dieckmann and Law, 1996). By computing the second and the cross derivative of the relative fitness, we can further demonstrate that, in this case, a singular strategies can only be convergent and non-invasible (CSS) or divergent and invasible (a repellor). Moreover, a CSS corresponds to a local maximum in emergent population properties, while a repellor corresponds to a local minimum, so that abundances, growth rates, and dynamical resilience always increase during evolution.

Figure 1 illustrates this study case for two generic intrinsic growth rate functions. Panels A and D show evolutionary trajectories, while panels B and E show corresponding pairwise invasibility plots (PIPs). Such plots show in gray, for a given resident phenotype on the x axis, the set of mutant phenotypes, axis y, that can invade it. They show the positions of evolutionary singularities (at the intersection of the diagonal and the zero fitness contour) and whether these strategies are convergent or divergent, invasible or non invasible (Geritz et al., 1998). For clarity, we report the direction of evolution determined from the PIP on panels C and F. Panels C and F show how the value of phenotype affects the emergent ecological properties. In both examples, evolution optimizes all emergent properties. Note that while this result is verified when singular strategies exist (panels A to C), it is also true when no evolutionary singular strategy exists and continuous directional selection occurs (panels D to F). General computations of the evolutionary singular strategies and their convergence and invasibility properties are given in the Online appendix A.

**Figure 1:**
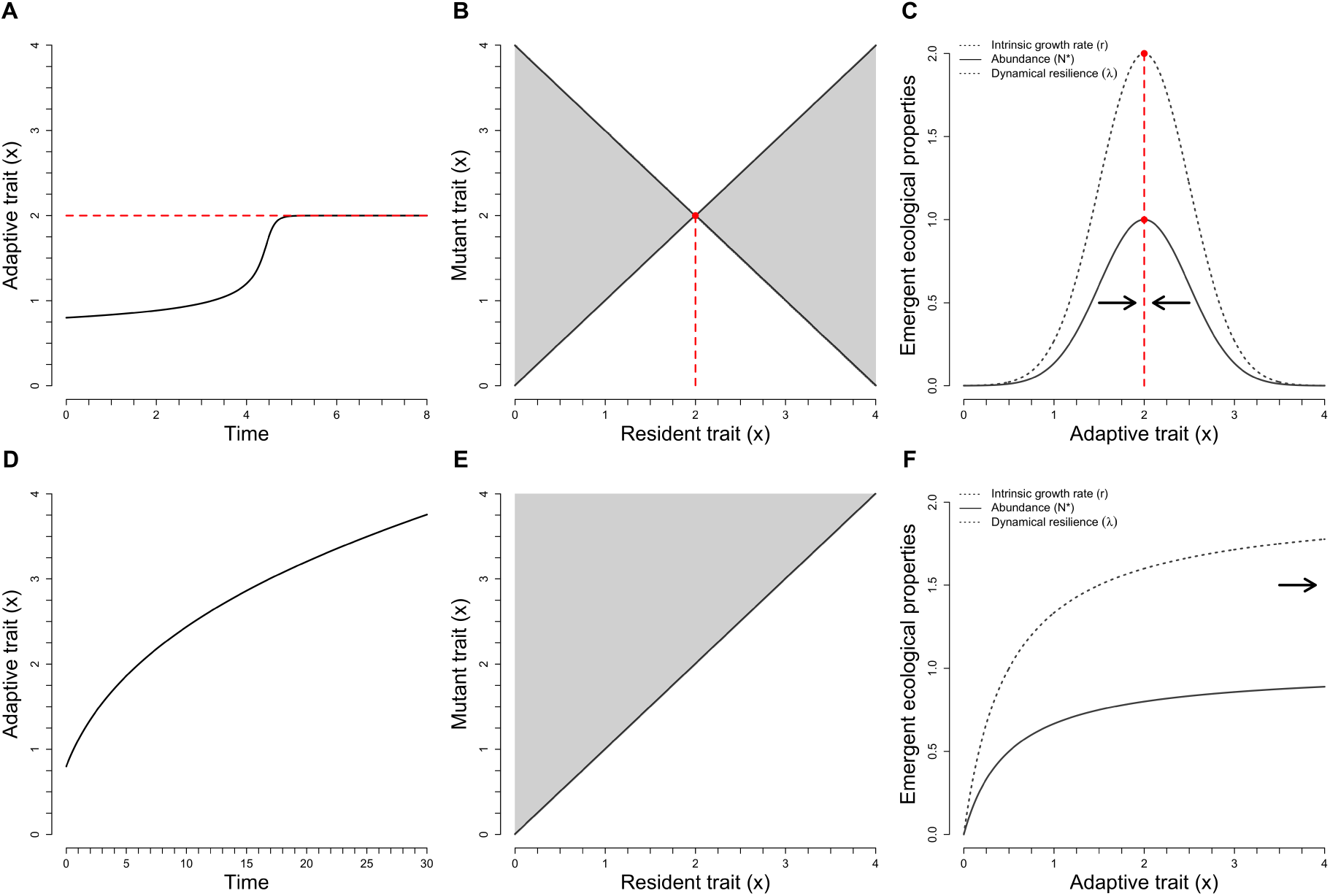
Two examples of eco-evolutionary dynamics leading to the optimization of intrinsic growth rates, abundances, and dynamical resilience. Panels A and D show the evolutionary trajectories, while panels B and E show corresponding PIP plots. Panels C and F show how the three emergent ecological properties change along phenotypic variations. Evolutionary singular strategies are in red and the black horizontal arrows show the direction of evolutionary trajectories. Panels A to C assume that growth rates are optimal when the phenotype *x* is at 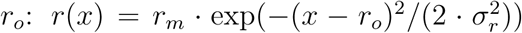, and the parameters value were set to *r*_*m*_ = 2, *r*_*o*_ = 2, *σ*_*r*_ = 1/2. Panels D to F assume growth rates to be a monotonically increasing and saturating function of the phenotype *r*(*x*) = *r*_*m*_ · *x*/(*h*_*r*_ + *x*), with parameters set to *r*_*m*_ = 2, *h*_*r*_ = 0.5, and *α* = 2

We now discuss what biological traits may fit this scenario. Here, the phenotype only affects the intrinsic growth rate of the population *r*(*x*). Recall that the intrinsic growth rate is simply *r*(*x*) = *f* (*x*) − *m*(*x*), with *f* (*x*) and *m*(*x*) the *per capita* fecundity and mortality rate respectively. It follows that, for a biological trait to fit this model, it should only impact the individual reproduction and survival, without impacting competitive interactions with other individuals of the population, as *α* is independent of the phenotype. We believe some biological traits may fit these criteria:

- Morphology or physiology of reproduction. For instance, many examples exist of assemblages of cryptic species, that seem ecologically equivalent and morphologically very similar, yet are well separated from a reproductive point of view (see McPeek and Gomulkiewicz (2005) for a review of different examples). It is generally assumed that these species are separated by key differences in reproductive traits only and maybe considered good examples of a diversity maintained by neutral processes (Leibold and McPeek, 2006), meaning that phenotypic differences are not supposed to affect competition. Evolution of such traits within populations likely affects fecundity rates (by constraining reproduction among individuals) while having no direct on competition, hence fitting the requirements of this class I model.
- Evolution of traits that are mostly constrained by fecundity vs survival trade-offs. Indeed, evolution of such traits affect the two components of *r*(*x*), while not directly changing competition among individuals. For instance, it has been proposed that some plant defense strategies (but not all, see (Strauss et al., 2002)) are costly in terms of growth (Herms and Mattson, 1992). More defended plant then have larger survival (i.e., lower *m*(*x*)) and lower fecundity (lower *f* (*x*)). As another example, consider the work of Rose et al. (2009). In this experimental study, plant fast growing phenotypes show greater sensitivity to external stress, so that survival is reduced, again fitting the requirement of our class I model.

### Optimization of abundance only

As a second study case, we consider that both the intrinsic growth rate, *r*(*x*) > 0, and intraspecific competition *α*(*x*) > 0 of Verhulst’s model depend on phenotype *x*. In such a situation, the invasion fitness a rare mutant *x*_*m*_ in a resident population is given by

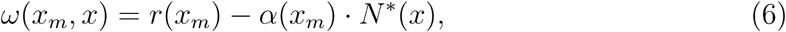

which is clearly again a class I fitness function. Optimization of species abundance therefore applies. However, as shown on the examples of figure 2, evolution no longer optimizes the intrinsic growth rate and the dynamical resilience. This can been shown by computing the fitness gradient:

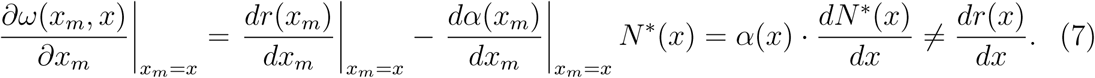

**Figure 2:**
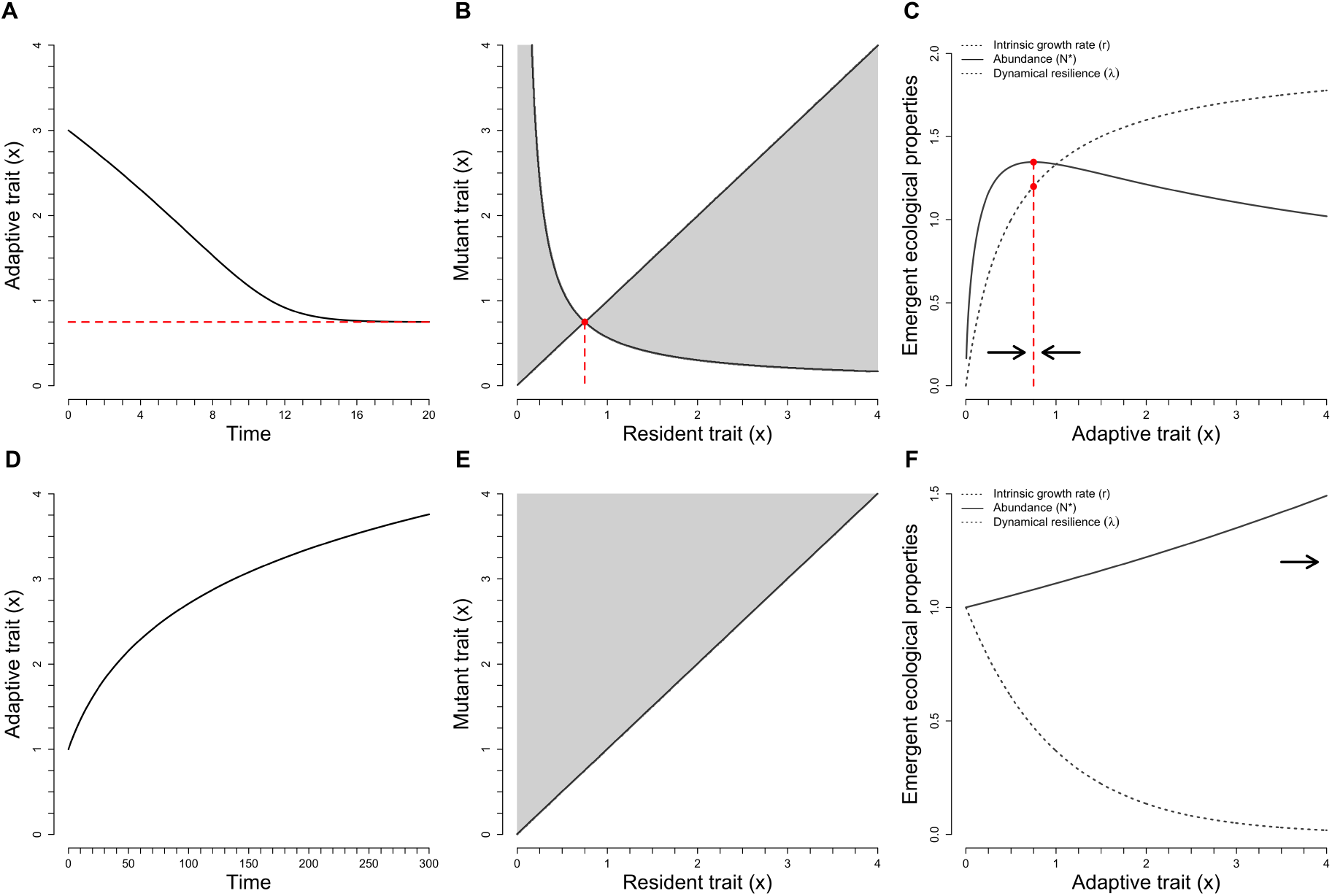
Two examples of eco-evolutionary dynamics leading to the optimization of abundances only. Panels A and D show the evolutionary trajectories, while panels B and E show the corresponding PIP plots. Panels C and F show how the three emergent ecological properties change along phenotypic variations. Evolutionary singular strategies are in red and the black horizontal arrows show the direction of evolutionary trajectories. Panels A to C assume the following generic functions *r*(*x*) = *r*_*m*_ · *x*/(*h*_*r*_ + *x*) and *α*(*x*) = *α*_0_ *· x*^*δ*^, with parameters set to *r*_*m*_ = 2, *h*_*r*_ = 1/2, *α*_*o*_ = 1, and *δ* = 0.4. Panels D to F assume *r*(*x*) = *r*_*m*_ · exp(−*λ*_*r*_*x*) and *α*(*x*) = *α*_*m*_·exp(−*λ*_*α*_*x*), with the following parameters: *r*_*m*_ = 1, *λ*_*r*_ = 1, *α*_*m*_ = 0 and *λ*_*α*_ = 1.1

Remembering that singular strategies require the fitness gradient to be zero, evolution optimizes population abundance, but in general one should not expect optimization of the intrinsic growth rate an neither of the the dynamical resilience.

Figure 2 illustrates this case for two sets of generic functions for *r*(*x*) and *α*(*x*). As in figure 1, panels A and D show the evolutionary trajectories, panels B and E show the PIP plot, and panels C and F show how evolutionary dynamics affect the various ecological properties we consider here. Figure 2 perfectly illustrates the optimization of abundance, but not of the intrinsic growth rate and of dynamical resilience. Note that while this result is verified when singular strategies exist (panels A to C), it is also true when no evolutionary singular strategy exists and continuous directional selection occurs (panels D to F). Details on the computation of evolutionary singular strategies, their convergence and invasibility properties are given in the Online appendix A.

Biological situations that correspond to this second model require that the phenotypic trait affects one or both components of the intrinsic growth rate (fecundity and/or survival) and the competitive ability of the species. Several examples of traits have been proposed that seem to fulfill these requirements:

- Returning to plant defenses, while some of these defenses mostly divert energy from intrinsic growth rate, as previously discussed, other studies suggest that other defenses directly impact the competitive ability of the plant. In such instances, survival is higher for highly defended strategies (enhancing *r*(*x*)) while their competitive ability is depressed. For instance, Agrawal et al. (2012) describe a field experiment on plant species *Oenothera biennis* in which suppression of seed predators leads to the fact counter-selection of defenses, with direct impacts for the competitive ability of the plant. Next to this particular example, this idea that some plant defenses are constrained by variations in competitive ability has been proposed to explain several cases of plant invasions (the “Evolution of Increased Competitive Ability” hypothesis (EICA) Blossey and Notzold (1995)). Invasive species, having fewer enemies in their new environment, then selects lower defenses and higher competitive abilities, which accelerates their invasion dynamics.
- More generally, one of the key axis of understanding of life history strategies is the *r* vs *K* trade-off selection theory (MacArthur and Wilson, 1967; Pianka, 1970), where fast growing strategies have low competitive abilities and are progressively replaced by more competitive (but slower growing) species during successions. For phenotypic traits that fall along this trade-off axis, we expect the results of the present model to apply.

### Non-optimization

As a third study case, we consider that the competition strength between mutants and residents now depends on both traits *x* and *x*_*m*_, i.e., *α*(*x*_*m*_, *x*) for the competitive effect of a resident *x* on a mutant *x*_*m*_. Therefore, the invasion fitness is given by

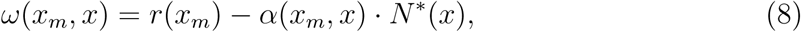

which is a class II fitness function. Because the resident is assumed to be at equilibrium *N*\*(*x*) = *r*(*x*)/*α*(*x, x*), the fitness gradient is given by

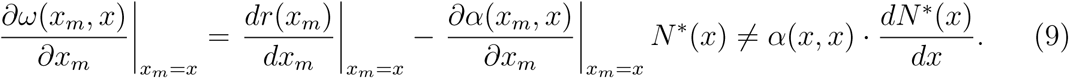

Contrary to the first two study cases (eqs. 5 and 7), the fitness gradient is not usually proportional to the gradient of abundance or of the intrinsic growth rate. Therefore, in general, evolution does not lead to the optimization of abundances or intrinsic growth rates. Also note that contrary to the previous cases, we do not necessarily get convergence to a fixed strategy (i.e., a CSS) or evolution diverging from the singularity (i.e., a repellor). Instead, all types of singular strategies are possible. Importantly, it is possible to get evolutionary strategies that are convergent but invasible. At such “branching points”, disruptive selection leads to the maintenance of polymorphism (see figs 3 panels D to F, 4, and 5 for an example). In such cases, trait displacement occurs among individuals of the population. Such a divergence leads to a decrease in competition of the individuals showing contrasted phenotypes (limiting similarity, see MacArthur and Levins (1967)).

**Figure 3:**
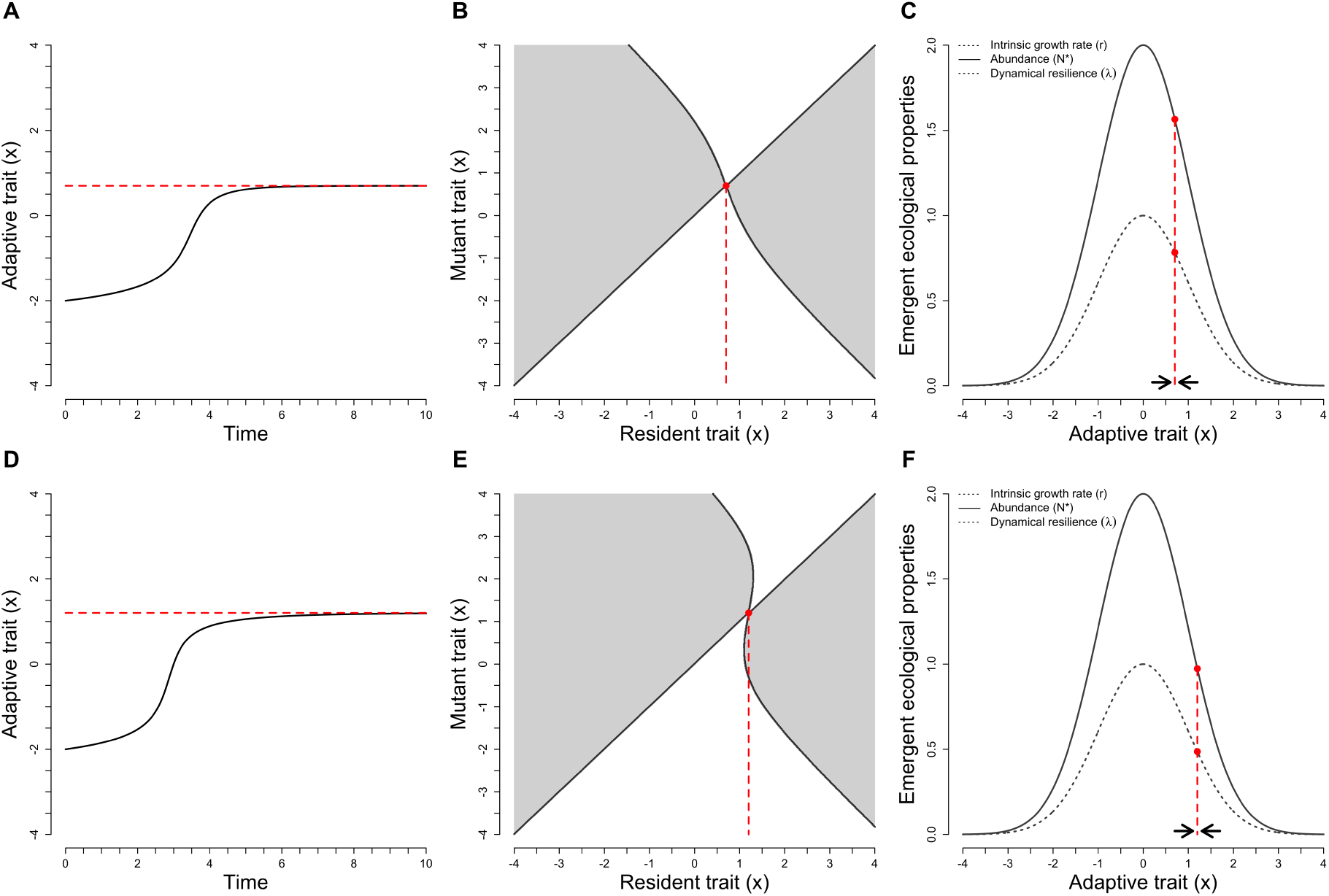
Two examples of eco-evolutionary dynamics illustrating that relative fitness functions of class II lead, in general, to the non-optimization of abundances, intrinsic growth rate, and dynamical resilience. Panels A and D show the evolutionary trajectories, while panels B and E show the corresponding PIP plots. Panels C and F show how the three emergent ecological properties change along phenotypic variations. Evolutionary singular strategies are in red and the black horizontal arrows show the direction of evolutionary trajectories. The competition model is given by: 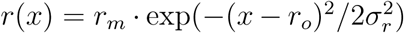 and the competition between two phenotypes *x* and *y* is given by *α*(*x, y*) = *c* ·(1 − 1/(1 + *ν* · exp(−*k* (*x y*)))), so that competition is asymmetric, favoring higher phenotypic values Kisdi (1999). The following parameters value were used: panels A to C with *r*_*m*_ = 1, *r*_*o*_ = 0, *σ*_*r*_ = 1, *c* = 1, *k* = 1.4, and *ν* = 1; and panels D to F with *r*_*m*_ = 1, *r*_*o*_ = 0, *σ*_*r*_ = 1, *c* = 1, *k* = 2.4, and *ν* = 1

Figure 3 illustrates this case, assuming that phenotypes compete, asymmetrically, along a gradient of resources as in Kisdi (1999). Depending on parameter values, the system can reach CSS (panels A to C) or an evolutionary branching strategy (panels D and F) (see also Kisdi (1999)). Consistent with the previous analysis (equ. 9), none of the ecological properties are maximized. Computations of evolutionary singular strategies, their convergence and invasibility properties are given in Kisdi (1999). After the branching (panels D to F), niche differentiation between two coevolving morphs will appear and the system eventually settle at a coalition of two CSSs (figs 4 and 5, see also Kisdi (1999)). This will be studied in section *Polymorphism coevolution after branching*.

**Figure 4:**
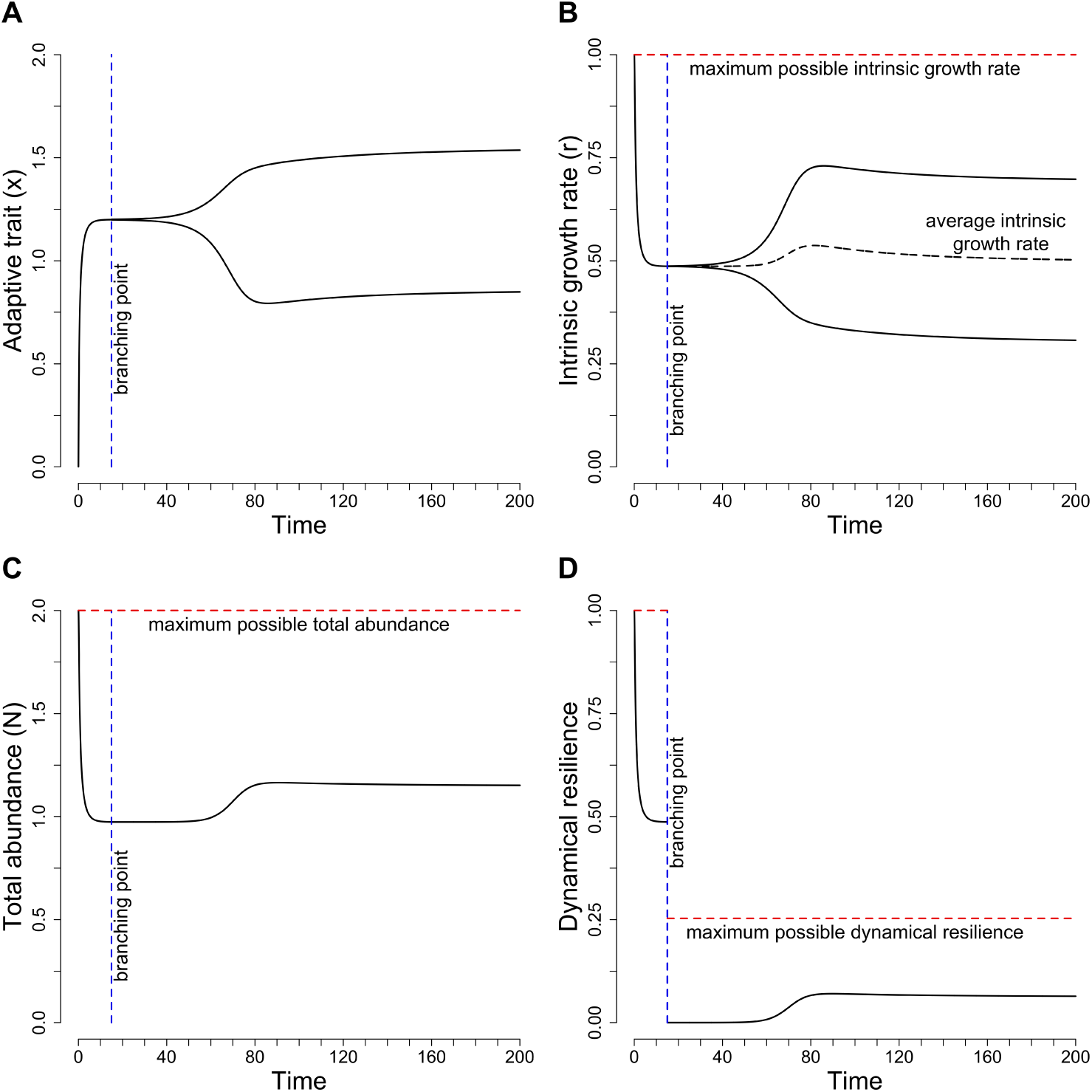
Co-evolutionary trajectories of two morphs after the branching point of figure 3 panels D to F. Panel A show the evolution of the phenotypic traits, it is the continuation of figure 4 panel D. Panel B to D show the evolution of the intrinsic growth rates, the total abundance, and the dynamical resilience, respectively. On all panels, the vertical blue dashed lines represent the time of the branching process. On panels B to D, the horizontal red dashed lines give the potential maximum value of the corresponding emergent properties, i.e., the maximum value that could be reached if one could chose the value of the phenotypes. On panel B, the black dashed line represents the evolution of the average intrinsic growth rate.

**Figure 5:**
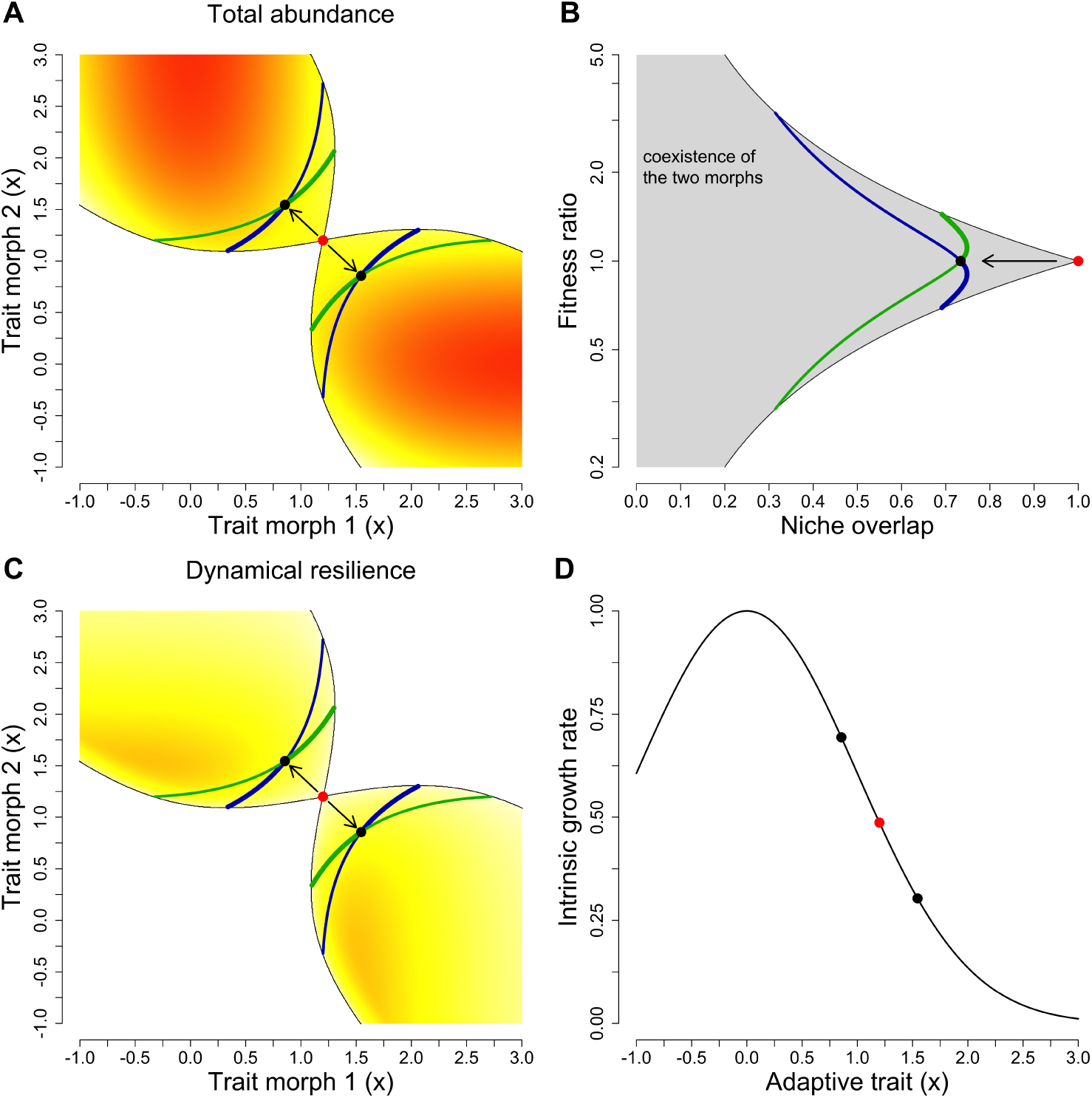
Co-evolution of the two morphs after the branching point of figure 3 panels D to F (see also figure for the corresponding branching process in time 4). The red dot shows the branching point, while the black dots shows the coalition of CSSs to which the system converges. Panels A and C show the traits evolution plots. The colored area corresponds to the protected dimorphic area. The color gradient on panel A gives the total abundance of the two morphs at ecological equilibrium, while on panel C it stands for the dynamical resilience. Panel B shows changes in the fitness-ratio (*κ*) and niche-overlap (*ρ*) for the two morphs during the branching process. Green and blue lines correspond to the evolutionary isoclines, i.e. the set of points at which evolutionary gradients equal zero for morph 1 and of morph 2, respectively. These two isoclines can be either invasible or non-invasible, depending on the sign of the second derivatives of the relative fitness functions. Thick lines show non-invasible strategies, while thin lines indicate invasibility. Panel D shows how coevolution affects intrinsic growth rates during the branching process.

We believe that many biological traits will fit this third model. More specifically:

- Although competitive ability can be presented as a characteristic of a given phenotype (as discussed earlier), the outcome of competition is most often dependent not only on the individual traits, but also on other traits present in the population. Considering competition for light as an example, plant size *x* may be competitive or not depending on the size *y* of individuals around, so that the outcome of competition will depend on *x* − *y* (see for instance Falster et al. (2017)).
- Now consider one of the most studied trait in ecology: body size (often measured as body mass). It is well known that variations in body mass likely affects intrinsic growth rates, by changing fecundity and mortality rates. For instance, Barneche et al. (2018) show that in fish species reproductive investment sharply increases with body size within species, so that large phenotypes have a very large impact on the population productivity. Variations in survival and in the whole intrinsic growth rate with body size are also well documented (Savage et al., 2004). Based on these components, consequences of body size evolution would follow class I models we introduced earlier. We expect this to be true in some situations. However, for some species, interference competition occurs and larger individuals often have a systematic advantage, while smaller individuals are often favored in scramble competition (Persson, 1985). In any of these cases, *α* will depend not only on the individual body size, but also on the trait of the interactor, fitting the current model. Similarly, predators of similar body sizes often have similar preys so that competition can be enhanced if they are close in body size (Woodward and Hildrew, 2002). While we so far presented *α* as a competition coefficient, note that it actually embodies any negative density dependent effect, including cannibalism. Again, cannibalism of individual of body size *x* by an individual of body size *y* depends on the relative difference in body size (Persson et al., 2003). For all these reasons, we expect that in situations where body size evolves fast, the current model will most often apply, so that no optimization can be expected. This is particularly important, as fast variations in body size are expected under current global changes, smaller sizes being usually favored in warmer environments (Daufresne et al., 2009) and when species are heavily harvested (Olsen et al., 2004; Conover and Munch, 2002).
- Another current interest given climate modifications is the possible evolution of phenologies. Such evolutionary changes have been observed in many species of different groups (e.g., Phillimore et al. (2010); Franks et al. (2007); Nussey et al. (2005)). Consider two phenotypes whose peak dates of activity in the year are *x* and *y* respectively. If these two phenotypes consume the same resources, competition for these resources will not depend on *x* or *y* only, but will be greater when phenologies overlap more. It will therefore depend on the matching between *x* and *y*, a situation that fits the current model.

## Ecological interpretation of the two classes of relative fitness function

There is a fundamental ecological difference between the two first study cases and the third one. The first two study cases assume no niche differentiation between the resident and the mutant phenotypes, while the third study case assumes that their interaction rely on niche differences, as competition is defined by the traits of the two interacting individuals. More generally, this is also true for the two classes of relative fitness function: the first class implicitly assumes no niche differentiation, while the second class does. From an ecological perspective, the dynamic resident-mutant can be considered as a special case of classical models of two-competing species. The outcome of such dynamics can therefore be studied within the framework of the modern coexistence theory, by computing the niche overlap of mutants and residents and their fitness ratio. Recall that in this framework, for a given fitness ratio, the species whose fitness is lower can only coexist with the other species if the overlap is below a threshold value (Chesson, 1990; Saavedra et al., 2017).

For the resident-mutant dynamic in study case 2, the niche overlap equals

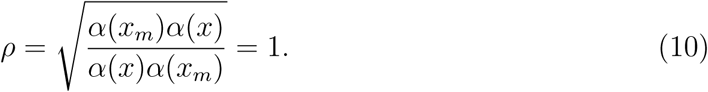

Remembering that in this study case the competition strength is a function of the resident traits *x*, or the mutant traits *x*_*m*_ but never of both traits (contrary to the third study case). In such situation, Niche difference (1 − *ρ*) then tends toward zero so that there is no room for coexistence between the resident and the mutant, one excluding the other. The outcome of the competition depends on the fitness ratio, computed by

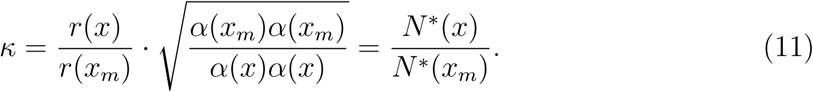

If *κ* > 1 the resident outcompetes the mutant and the mutant cannot invade, while if *κ* < 1 the mutant replaces the resident. As *κ* is, in this case, given by that abundance ratio, it follows that evolution optimizes abundance. When intraspecific competition *α* does not depend on phenotype *x*, which is our study case 1, *κ* simplifies to the intrinsic growth rate ratio *κ* = *r*(*x*)/*r*(*x*_*m*_), so that evolution also maximizes intrinsic growth rates.

In our third study case, the niche overlap can be computed as

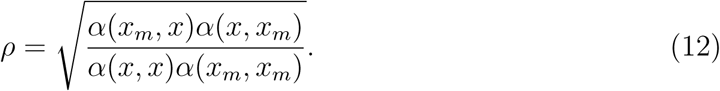

In general *ρ* ≠ 1. When *ρ* < 1, there is room for coexistence between resident *x* and mutant *x*_*m*_. Now the fitness difference reads as

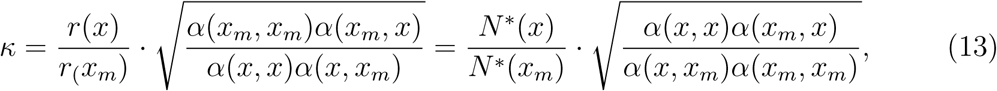

and coexistence of the two phenotypes requires that 1/*ρ* > *κ* > *ρ*. If the first inequality is violated, the resident wins and the mutant cannot invade, while if second inequality is violated the mutant replaces the resident. Contrary the the previous cases, evolution will not necessarily increase abundance or intrinsic growth rates, as selection of larger abundance can be counter-balanced by the competitive imbalance term

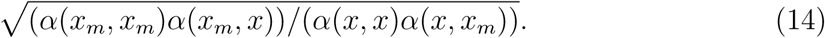

Importantly, coexistence conditions between the resident and the mutant can be fulfilled, so that a branching becomes possible.

## Proof of the main result

We will give the general idea behind the proof of our main results. The full mathematical proof is given in Appendix B. We start by showing that class I of models leads to evolution that optimizes abundance and to evolutionary singular strategies that can only be CSS or reppelor. Then we will explain why for class II models, such optimization principles do not generally hold, and why any type of singular strategies can be expected.

The first step of our proof consists in linking the evolutionary gradient with the gradient in abundance. For the class I models, the ecological equilibrium *N*\*(*x*) of a population of traits *x* must fulfilled *f* (*x, N*\*(*x*)) = 0. By taking the total derivative relative to the trait *x* we obtain the following equivalence

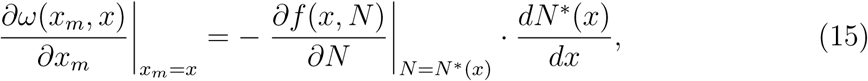

It generalizes equations (5) and (7) of the first and second study cases, respectively. Therefore, an evolutionary singular strategy, i.e., a trait value *x*\* at which the fitness gradient vanishes, is equivalent to a local optimum in abundance, i.e., a trait value at which the derivative of the abundance vanishes. This proves, for relative fitness function of class I, that an evolutionary singular strategy is equivalent to a local optimum (maximum or minimum) in abundance.

Linking the property of the optimum (if it is a maximum or a minimum) with the type of singular strategies (convergent or divergent and invasible or non-invisible) needs to study the second and the cross derivative of the relative fitness function (Dieckmann and Law, 1996). The details are given in the Appendix B. The outline of this part of the proof is the following. We start by showing that the cross derivative of the relative fitness function always vanishes. This demonstrates that the singular strategy must by convergent and non-invasible (CSS) or divergent and invasible (reppelor). Then we prove that the second derivative of the relative fitness function is positively proportional to the second derivative of the abundance at equilibrium. This demonstrates that a CSS corresponds to a local maximum, while a reppelor is equivalent to a local minimum.

Now, we explain why for class II models, such an optimization principle does not hold. Again, at the ecological equilibrium *N*\*(*x*) we have must have *f* (*x, x, N*\*(*x*)) = 0. By taking its derivative relative to *x*, we obtain this time

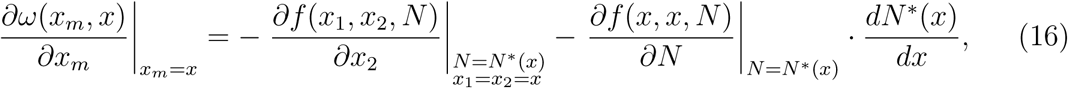

a more complex equation than the one in the simplest case (equ. 15). The partial derivative relative to *x*_2_ (first term on the left-hand side) does not necessarily equals zero, so that an evolutionary singularity is no longer equivalent to an optimum in abundance. Furthermore, we can proof that the cross derivative of the relative fitness function does not anymore vanishes in general. Consequently, any type of evolutionary singular strategy is possible.

## Some remark

First remark: the two first study cases, which optimize intrinsic growth rate and/or abundance, can be related to the recent results by Metz et al. Metz et al. (2008); Metz and Geritz (2016) on optimization. Actually the relative fitness of our first study case (equ. 1), can be rewritten as *ω*(*x*_*m*_, *x*) = *r*(*x*_*m*_) − *r*_(_*x*) = *α* · (*N*\*(*x*_*m*_) − *N*(*x*)), which implies optimization of *r* and *N*\* according to the corollary 3.4 of Metz and Geritz (2016). The second study case can be rewritten as *ω*(*x*_*m*_, *x*) = *α*(*x*_*m*_) *·* (*N*\*(*x*_*m*_) − *N*(*x*)), which implies optimization of *N*\* only. The first study case can be rewritten, more generally, as *ω*(*x, x*_*m*_) = *r*(*x*_*m*_) − *ϕ*(*N*\*(*x*)), where *r*(*x*_*m*_) > 0 is the intrinsic growth rate of the mutant and *ϕ*(*N*\*(*x*)) is the effect of the resident population *x* on the per capita growth rate of the mutant. Assuming *ϕ* being a monotonically growing function, which is equivalent to a negative density dependence (intraspecific competition), one can demonstrate that evolution optimizes all three properties (see also example 6.1 of Metz et al. (2008)).

Second remark: the two class of relative functions can be multiplied by any positive function *ψ*(*x*_*m*_, *x*) > 0 without changing the results on optimality, and on convergence and invasibility (see also (Metz et al., 2008; Metz and Geritz, 2016; Lion and Metz, 2018)).

## Polymorphism coevolution after branching

Figure 3, panels D to F show that for class II models, a branching point is possible. After this branching point, two morphs appear and coevolve. Figures 4 and 5 shows this polymorphism coevolution. From an ecological perspective, the dynamic of the phenotypes can be constructed as a special case of a two competing species model, given by the following set of equations

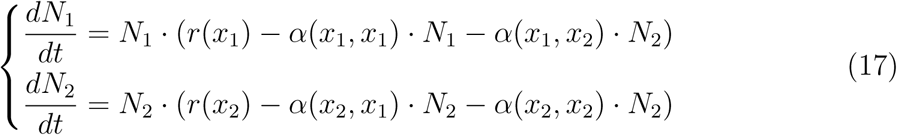

The intrinsic growth rate *r*(*x*_*i*_) and interaction strength *α*(*x*_*i*_, *x*_*j*_) are functions of the morphs traits *x*_1_ and *x*_2_. Figure 4 show the evolution as function of the time. In particular, it shows the evolution of the phenotypic trait value and the three emergent properties considered here. On figure 5 panels A and C, the colored areas define the so-called protected dimorphism area. Ecologically, it correspond to pairs of phenotypic traits such that the two morphs coexist, i.e., equation (17) leads to a positive and stable equilibrium point 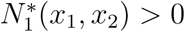 and 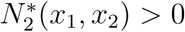 (Kisdi, 1999). In coexistence theory this region can be represented in the fitness-ratio (*κ*) and niche-overlap (*ρ*) space. Specifically, these two metrics are computed as

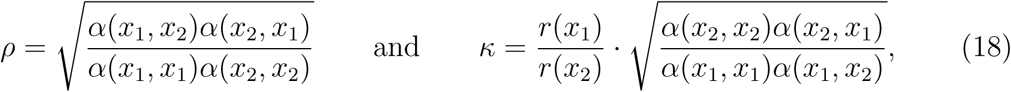

and coexistence is reached when 1/*ρ* > *κ* > *ρ*. These two inequalities are shown on panel B (fig. 5). Note that if interaction strengths are not a function of the morph traits (*α*) or a function of only the first argument *α*(*x*_*i*_), the niche overlap equals one. Coexistence between the two morphs is then impossible and the protected dimorphic area vanishes.

Figures 4 and 5 show, as expected, that the branching dynamics reduce the overall level of competition, as the niches of the two morphs diverge, so that total abundance increases. Note however that abundances do not reach the potential maximum. This increase is coherent with figure 5 panel B showing that the coevolution of the two morphs decreases their niche overlap. Interestingly, the two morphs remain equivalent from a fitness point of view (*κ* = 1, fig. 5 panel B). Panels C and D show that coevolution neither optimizes dynamical resilience nor intrinsic growth rates.

Our main results on optimization of emergent properties can be generalized to polymorphic coevolution or to species coevolution as follow. The evolution of each morph *i* (or species) is defined by its relative fitness function *ω*_*i*_(*x*_*i,m*_, ***x***), i.e., the per capita growth rate of a rare mutant of trait *x*_*i,m*_ in a resident community of *n* morphs (or species) of traits ***x*** = (*x*_1_, *x*_2_, …, *x*_*n*_) at ecological equilibrium 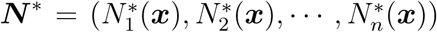. Our two classes of relative fitness function generalize as follow

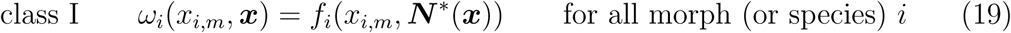

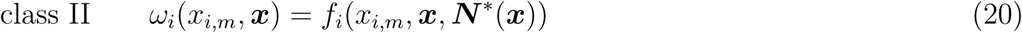

In Appendix C, we generalize our proof to the coevolution case. We demonstrate that class I fitness functions lead to coevolution that optimize abundances and to singular strategies that are convergent and non-invasible (CSS) or not-convergent and invasible (repellor). While class II fitness functions lead to coevolution that, in general, do not optimize abundance and that any type of singular strategies is possible. As in the monomorphic case, we believe that many biological situations will fall in the second class. It is also the second class that may allow the emergence of polymorphisms through branching events and the corresponding niche differentiation.

## Discussion

Understanding whether evolution leads to optimality in intrinsic growth rate, abundance, or dynamical resilience is not only a theoretical question, but has practical implications for conservation, for the sustainable management of exploited species, and for invasion biology among others. After a theoretical discussion, we will discuss these empirical implications.

Whether evolution optimizes *r* (or *R*_0_) and/or abundance is a long standing question. The belief that evolution maximizes those population properties, i.e., evolution happens for the good of the species is still widespread. Dieckmann and Ferrière (2004) argued that this dogma arises by not considering frequency and density dependent selection. Indeed, quite often intrinsic growth rate (or one of its component) or variations in abundance are considered as good proxies to assess the fitness landscape. This, however, ignores the fact that this landscape is dynamic, due to frequency or density dependent selection. When considering frequency and density dependent fitness evolution becomes much richer than a simple optimization process (branching point, evolutionary suicides or murders, among others). That being said, it is not enough to introduce solely frequency and density selection to stay away from *r* and/or abundance optimization. As shown above, but also in Metz et al. (2008) or Lion and Metz (2018), optimization of *r* (or *R*_0_) can still be achieved for specific form of invasion fitness (one-dimensional environment feedback loop).

In this manuscript, we went a step forward by proposing a classification of invasion fitness into two generic classes (eqs. 1 and 2). The difference between our two classes may look very subtle; class II assumes that the invasion fitness depends directly of the resident and the mutant trait, while in class I it is assumed that the resident trait acts only through the density of the resident population. However, this subtle distinction leads to opposite results on optimality, convergence and invasibility properties, and niche displacement. Class I systematically selects phenotypes with larger abundance, that is, evolution follows a *K*-selection process, while, in general, class II does not exhibit such a *K*-selection. We also demonstrate that evolutionary branching is impossible in class I, while it is possible in class II. We explained that this is intimately linked to niche displacement among phenotypes. In class I, there is no possibility for niche differentiation, in the sense of coexistence theory, between the mutant and the resident phenotypes, while class II can lead to niche differentiation. Putting these results together, they suggest that we have either

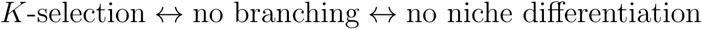

or

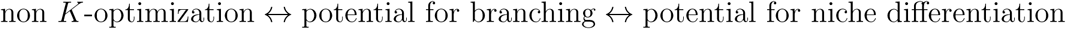

The latest result, on niche displacement, arise when studying the ecological dynamics between resident and mutant phenotypes under the lens of coexistence theory (Chesson, 1990; Saavedra et al., 2017). Class I invasion fitness functions lead to ecological dynamics that do not allow niche differentiation between mutant and resident phenotypes, thereby preventing their coexistence. This implies that evolutionary dynamics cannot exhibit branching events, so that no polymorphism emerge in class I. Intuitively this results from the fact that the interaction effect of a mutant on another mutant and the effect of a resident on a mutant are similar, because the relative fitness function is a function of only the mutant traits and the resident abundance. Fitness ratio metrics then show that evolution will systematically select the morph with the largest abundance (*K*-selection). We could further constrain our class I and reach the *r* optimization of Metz et al. (2008). Class II functions allow niche differentiation, because the invasion fitness is a function of both the mutant and the resident traits. Residents and mutants can then coexist, which ultimately may lead to branching and polymorphism emergence. However, using fitness ratio, we showed that then evolution does not select necessarily larger abundance, because niche difference implies in general competitive imbalance which can counterselect morphs with larger *r* or abundance. This result is a step toward a better understanding on how ecological niche construction relates to evolution (Odling-Smee et al., 2003; Lewontin, 2013).

Our results suggest that evolution will seldom optimize population growth rates. Such optimization only happens for particular models within the first class. More specifically, it requires that the evolving trait only affects the intrinsic growth rate of the evolving species. While we propose this may occur for some traits (e.g., traits that are linked to the reproductive systems but do not directly modify ecological interactions, or traits whose trade-offs only involve intrinsic reproduction or mortality), we do not expect it to generally occur. This has important consequences for various management or conservation aspects. Concerning conservation, restoring intrinsic growth rates is a very important objective as it would allow an exponential growth of the rare populations. Evolution is now more widely considered in conservation plans (Stockwell et al., 2003). While our results suggest that natural selection will not necessarily help to maximize the growth rate, we still agree that evolution in general and genetic variability in particular should be carefully considered in concervation plans. Indeed, such restorations of variability would alleviate negative effects of genetic drift or inbreeding, aspects that are not explicitly accounted in our analysis, but likely very important in rare populations. Populations growth rate will then be increased, as detrimental mutations will be less easily fixed.

Management of intrinsic growth rates is equally important for invasive species. As these species typically start from a restricted set of individuals, their vast increase in population size or area occupied is in essence a problem of population growth rate. We believe that our results may have several implications in this regard. Note that fast evolution in various traits have been observed in invasive species (Mooney and Cleland, 2001). Whether such an evolution will further increase the growth rate of the species, thereby making possible side-effects of the invasion larger is an important matter. Our results suggest that we should not necessarily expect such an optimization. Particularly, defense traits have repeatedly evolved in invasive species (e.g., Müller-Schärer et al. (2004)), as these species are often freed from some of their natural enemies (“enemy release hypothersis”). Because such traits directly affect ecological interactions, we expect them to follow our second class model. Therefore their evolution should not optimize the invasive species intrinsic growth rate. A similar line of reasoning could be applied to epidemics. Avoiding epidemics largely relies on the control of the disease growth rate (that is, keeping the *R*_0_ below one). As disease agents often have fast generation times and large populations, evolution readily occurs and may affect this *R*_0_. Our approach suggests that, except for the most simplified ecological scenarios, such evolution will not optimize the *R*_0_ (see Lion and Metz (2018) for a more complete analysis).

Next to the management of species intrinsic growth rates, understanding the effects of evolution on population density or biomass is equally important. Most food resources consumed by human population stem from the use of exploited species, wild (e.g., fisheries) or domesticated (e.g., agriculture). In these different instances, the amount of resources is correlated to the density or biomass available. From an agricultural point of view, humans may guide the evolution of the species directly through artificial selection. Our results highlight that the relationship between the trait that is modified and the ecological interactions of the cultivated species with other species of the ecosystem will largely determine whether optimization can be expected or not. Because the interaction context is often simplified in selection assays, this highlights that many traits that are selected may eventually lead to non optimal population sizes in the field (see also (Loeuille et al., 2013)). Now consider wild species. While direct artificial selection is then not possible, harvesting these species incurs large extra-mortality. Such large selective pressures often yield fast evolution in the exploited species. For instance, in many fished species, evolution to earlier maturity at smaller adult size has been observed (Olsen et al., 2004; Grift et al., 2003), reviewed in (Edeline and Loeuille, 2020). Because body size is involved in many ecological interactions, we expect its evolution to belong to class II models. Therefore it is quite possible for this evolution to be detrimental to the standing density or biomass.

Our results also highlight that evolution may readily produce trade-offs among different emergent properties. Indeed, optimization of all three emergent population properties (growth rate, standing biomass and resilience) we studied only occurs in the simplest cases (ie, when the trait only affects the species intrinsic growth rate). In all other instances, any optimality in one of the emergent properties will come at a cost in one of the other. This has important applied consequences. Indeed, in many cases, ecological management is not simply about improving one population aspect, but several ones. Conservation of rare species often aim at improving their growth rates (see above), but also to maintain them above a given population threshold. In the case of fisheries, it is important not only to maintain the abundances (to get a certain yield but also to avoid unwanted ecological consequences such as population crash), but also to maintain the resilience of the system (e.g., Conover and Munch (2002)). Given evolution of maturity and body size in harvested species, we expect their evolution to follow class two models. Therefore we expect that such evolutionary dynamics are unlikely to foster a double objective of high abundances and large resilience. In line with this idea, note that the evolution of earlier maturity in Newfoundland cod stocks may be one of the reasons for the lack of recovery of the population, and has been directly linked to changes in resilience (Olsen et al., 2004).

To conclude, we propose that, while we have worked here on very simple models in very simplified situations, the topic at hand may have far reaching implications not only from a conceptual and theoretical point of view, but also affect how we view the conservation and management of natural systems. This is especially true given the fastly accumlating evidence that evolution occurs of quite short timescales (Hairston et al., 2005) especially given current global changes (Urban et al., 2016). Exact implications for more complex systems (e.g., in the case of diffuse coevolution in complex ecological networks) or for specific cases require further investigation, and will certainly bring a new set of exciting questions.

## Online Appendix A: details of studied cases models

In this appendix, we provide the mathematical details of the different examples of ecoevolutionary models we presented in the main text.

### Model of fig. 1 panels A to C

Ecological dynamics are given by

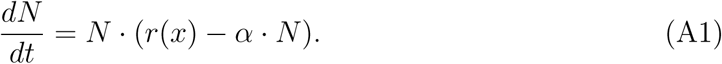

Only the intrinsic growth rate *r*(*x*) is function of the adaptive trait *x*. This function is given by a bell-shape function of the form

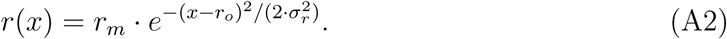

Parameters are *r*_*m*_ the maximum intrinsic growth rate, *r*_*o*_ the optimum value in the trait *x*, and *σ*_*r*_ the width of the bell-shape curve. Note that *r*(*x*) > 0 for all trait value *x*. Thus the ecological equilibrium is given by

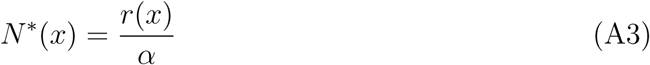

The relative fitness function of a rare mutant *x*_*m*_ in a resident population *x* is given by

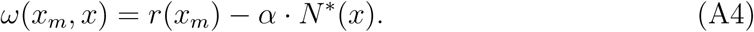

Then, the fitness gradient equals

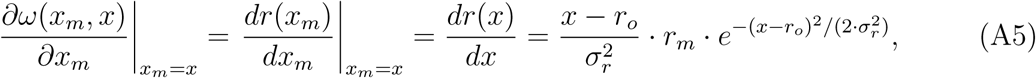

which equals zeros at *x*\* = *r*_*o*_. Evolutionary singular strategies match the optimum in intrinsic growth rate *x*\* = *r*_*o*_. As the ecological equilibrium equals *N*\*(*x*) = *r*(*x*)/*α*, thus *dN*\*(*x*)/*dx* = *dr*(*x*)/*dx ·* 1/*α*. This proves the equalities (5). Second and cross derivatives of the relative fitness function, evaluated at the singular strategy *x*\* are given by

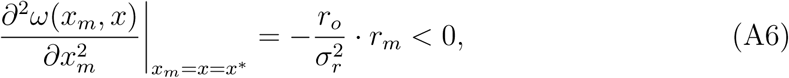

and

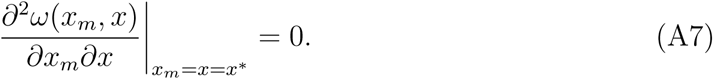

This proves that the singular strategy *x*\* = *r*_*o*_ is convergent and non invasible, i.e., a continuously stable strategy (CSS)

### Model of fig. 1 panels D to F

Ecological dynamics are given by

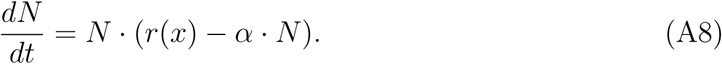

Only the intrinsic growth rate *r*(*x*) is function of the adaptive trait *x*. This function is given by a saturating function of the Michaelis–Menten form

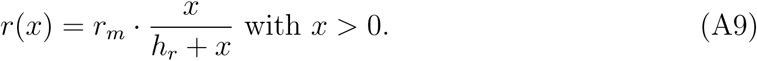

Parameters are *r*_*m*_ the maximum intrinsic growth rate and *h*_*r*_ the half saturation constant. Note that *r*(*x*) > 0 for all trait values *x* > 0. Thus the ecological equilibrium is given by

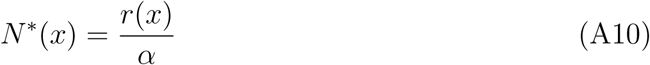

The relative fitness function of a rare mutant *x*_*m*_ in a resident population *x* is given by

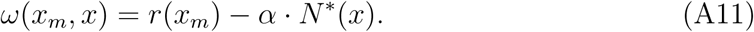

Then, the fitness gradient equals

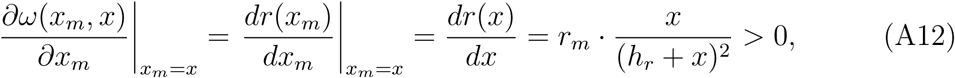

which is always positive. Selection therefore always favors larger *x* phenotypes, so that evolution also maximizes intrinsic growth rates and abundances. As the ecological equilibrium equals *N*\*(*x*) = *r*(*x*)/*α*, thus *dN*\*(*x*)/*dx* = *dr*(*x*)/*dx* · 1/*α*, this proves the equalities (5).

### Model of fig. 2 panels A to C

Ecological dynamics are given by

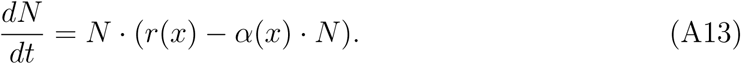

Both the intrinsic growth rate *r*(*x*) and the intra-specific competition are functions of the adaptive trait *x*. These two functions are given by

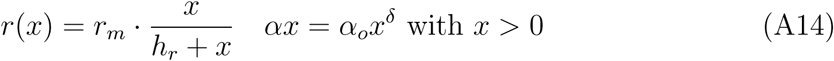

Parameters are *r*_*m*_ the maximum intrinsic growth rate and *h*_*r*_ the the half-saturation, and 0 < *δ* < 1 determines the shape of the trade-off function and *α*_*o*_ its amplitude. Note that *r*(*x*) > 0 for all trait value *x* > 0. Thus the ecological equilibrium is given by

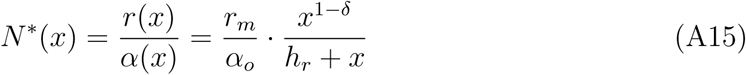

The relative fitness function of a rare mutant *x*_*m*_ in a resident population *x* is given by

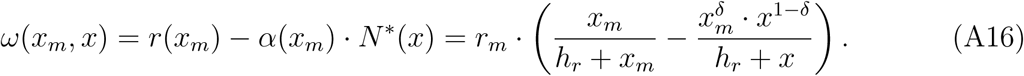

Then, the fitness gradient equals

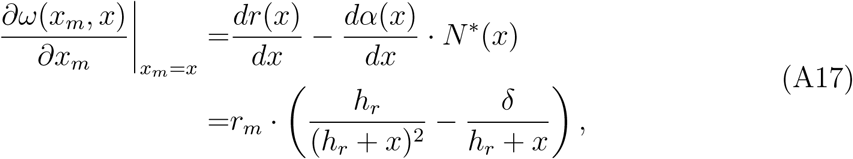

which equals zeros at *x*\* = *h ·* (1 − *δ*)/*δ*. Evolutionary singular strategies therefore do not maximize intrinsic growth rates, as the maximum of *r*(*x*) is reach when *x* → ∞. As the ecological equilibrium equals *N*\*(*x*) = *r*(*x*)/*α*(*x*), thus

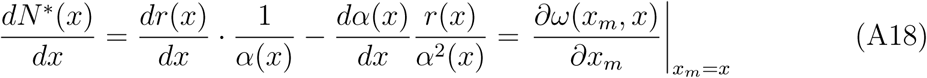

This prove the equalities (7). Furthermore the cross derivative of the relative fitness function, evaluated at the singular strategy *x*\* is given by

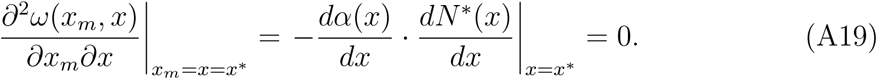

This proves that the singular strategy *x*\* = *r*_*o*_ can only be convergent and non invasible (CSS) or non convergent and invasible (reppelor).

### Model of fig. 2 panels D to F

Ecological dynamics are given by

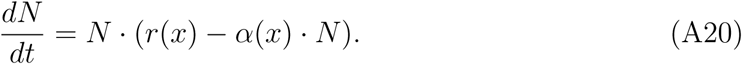

Both the intrinsic growth rate *r*(*x*) and the intra-specific competition are function of the adaptive trait *x*. These two functions are given by

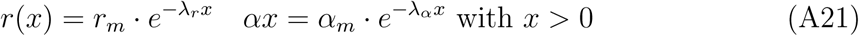

Parameters are *r*_*m*_ the maximum intrinsic growth rate, *α*_*m*_ the maximum level of intra-specific competition, *λ*_*r*_ > the rate at which the intrinsic growth rate decreases with the trait value *x*, and *λ*_*α*_ > the rate at which the intrinsic growth rate decreases with the trait value *x*. Note that *r*(*x*) > 0 for all trait value *x* > 0. Thus the ecological equilibrium is given by

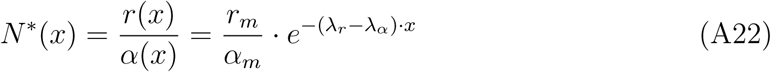

The relative fitness function of a rare mutant *x*_*m*_ in a resident population *x* is given by

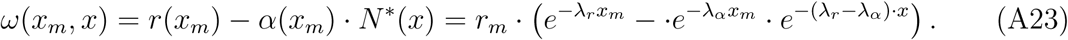

Then, the fitness gradient equals

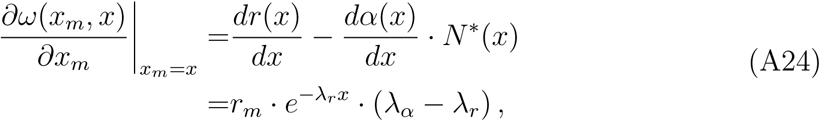

which is either positive or negative depending on the sign of *λ*_*α*_ − *λ*_*r*_. Thus phenotypic traits converge either to *x*\* = 0 or *x*\* → ∞. In the first case, abundance is maximized, while in the second case intrinsic growth rate is maximized.

## Online Appendix B: Proof of the main result

In this appendix, we give the mathematical details of the proof of our main results. We will start showing that class I models lead to evolution that optimizes abundance and to evolutionary singular strategies that can only be CSS or reppelor. Then we will explain why for class II models, such optimization principles do not generally hold, and why any type of singular strategies can be expected.

The proof works as follow. First let us assume that there exists a feasible ecological equilibrium *N*\*(*x*) > 0 that is function of the adaptive trait value. By definition of such an ecological equilibrium, the following equality must be fulfilled:

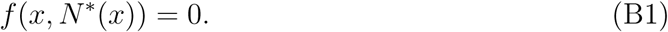

This is so, because *f* (*x, N*) is the *per capita* growth of the population given its abundance *N*, i.e., *dN*/*dt* = *N* · *f* (*x, N*). This equation leads to the relative fitness function of class I given by equation (1). By taking the derivative relative to *x* on both side of the above equation we obtain,

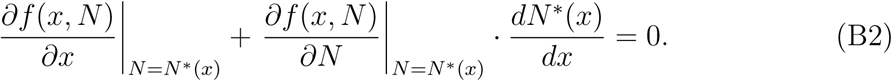

Now recall that, assuming small mutations, the fitness gradient is by definition:

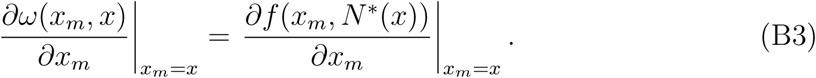

Combining these two equations results in

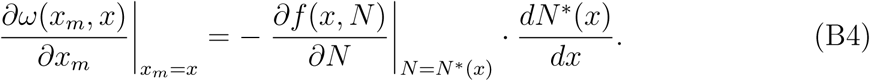

In general, we could expect *∂f* (*x, N*)/*∂N* |_*N*=*N*\*(*x*)_ < 0 as the model relies on negative density dependence (intraspecific competition). Therefore, an evolutionary singular strategy, i.e., a trait value *x*\* at which the fitness gradient vanishes, is equivalent to a local optimum in abundance, i.e., a trait value at which the derivative of the abundance vanishes.

This proves, for relative fitness function of class I, that an evolutionary singular strategy is equivalent to a local optimum (maximum or minimum) in abundance.

Evolutionary singular strategies can be either convergent (in which case local trajectories will converge to the strategy) or divergent (in which case natural selection favors strategies away from the singular strategy). They can also be invasible by nearby mutants, or non-invasible (ESS). Convergence and invasibility of singular strategies can be investigated using second derivatives of the relative fitness, assessed at the singularity (Dieckmann and Law, 1996). The singularity is convergent if and only if:

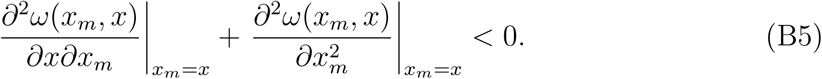

The singularity is non invasible if and only if:

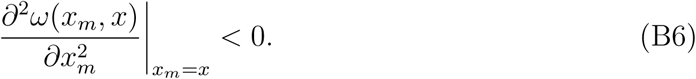

The cross derivative is given by

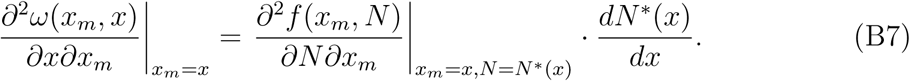

As an evolutionary singular strategy *x*\* corresponds to a local optimum in species abundance, the cross derivative vanishes at *x*\*. This proof that *x*\* can only be a CSS (convergent and non invasible) or a repellor (non-convergent and invasible), depending on the sign to the second derivative of the relative fitness function. A positive second derivative corresponds to a CSS, while a negative value results in a repellor.

Finally, we relate the second derivative of the relative fitness function to the second derivative of the ecological equilibrium. To do so, we take the second derivative, relative to the evolving traits *x*, of the equilibrium condition, *f* (*x, N*\*(*x*)) = 0. It leads to

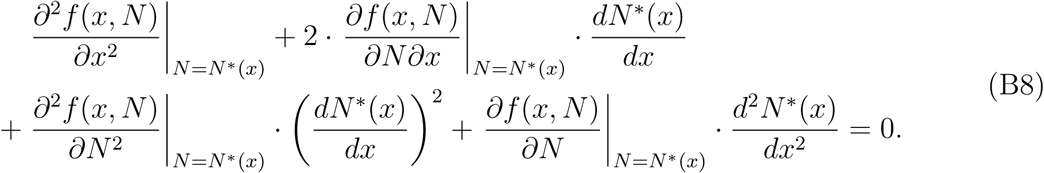

At an evolutionary singular point *x*\* the first derivative of the abundance function *N*\*(*x*) vanishes. Moreover, the first term equals the second derivative of the relative fitness function. Therefore, we obtain the following equivalence,

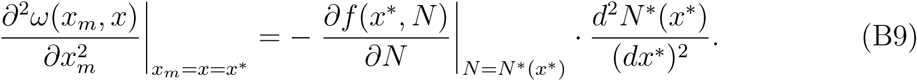

As advocated above, *∂f* (*x, N*)/*∂N* |_*N*=*N*\*(*x*)_ < 0, therefore the second derivative of the relative fitness function has the same sign as the second derivative of the abundance function. This ends the proof that a CSS leads to a local maximum in abundance, while a repellor results in local minimum and that an evolutionary singular strategy can only be CSS and repellor.

We now show that for class II models, such an optimization principle does not hold. Again, at the ecological equilibrium *N*\*(*x*) we have,

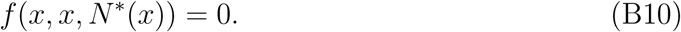

By taking its derivative relative to x, we obtain

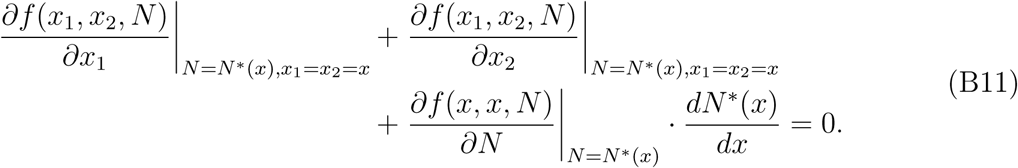

In the first term we can recognize the fitness gradient, which leads to

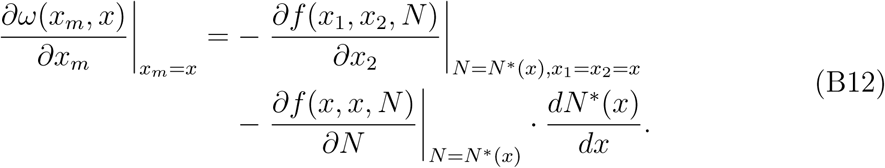

The term 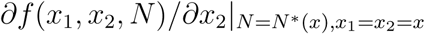 does not necessarily equals zero, so that an evolutionary singularity is no longer equivalent to an optimum in abundance. Moreover, the cross derivative of the relative fitness function, in class II, equals to

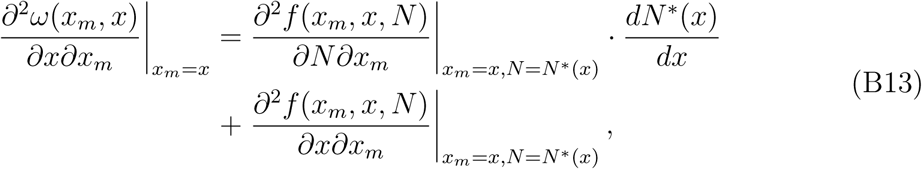

which, in general, does not vanish. This shows that, in general, evolution in class II models does not optimize abundance, and that any type of evolutionary singular strategy is possible.

## Online Appendix C: polymorphism or species coevolution

In this appendix, we give the mathematical proof of the generalization of our main results to polymorphism or species coevolution. We will start by class I fitness functions, for which the optimization principle hold, and then explain why for class II fitness functions such a principle does not hold in general.

### Class I of relative fitness function; optimization

Let us consider as set of *n* morphs (or species) whose ecological dynamics follow:

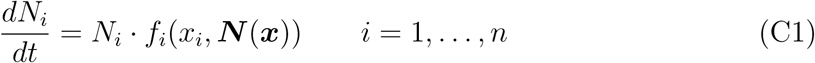

Moreover, we assume the existence of a stable a feasible ecological equilibrium ***N*** *(***x***) > 0, the solution of the following set of equations:

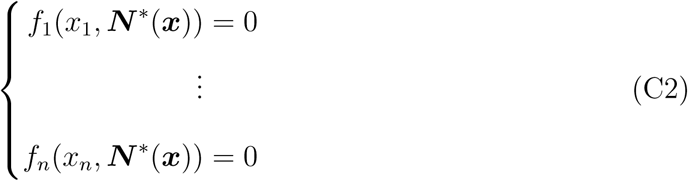

The relative fitness function of, and its derivative (the evolutionary gradient), for a mutant *x*_*i,m*_ of morph *i* are given by

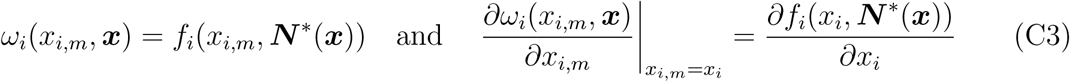

As in the monomorphic case, we can relate the abundance gradient to the evolutionary gradient in the following way. We take the total derivative of the set of equilibrium equations C2 relative to the traits *x*_*i*_ of morphs *i*. This results in the following set of linear equation for the abundance gradient *∂****N*** */*∂x*_*i*_:

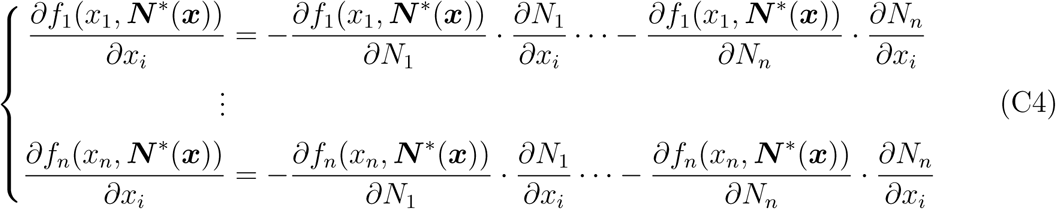

This set of linear equation can be rewritten in a matrix format as

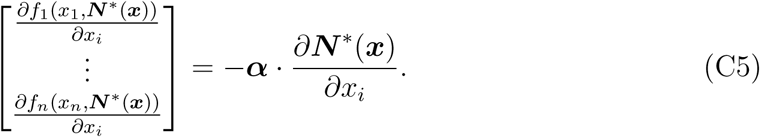

The elements of the matrix ***α*** are the partial derivatives *∂f*_*i*_(*x*_*i*_, ***N*** *(***x***))/*∂N*_*j*_ of the per capita growth rate relative to the abundance evaluated at the ecological equilibrium. Therefore the matrix ***α*** can be seen as the interaction matrix at ecological equilibrium. For class I fitness functions, all terms of the vector of the left term equals zeros with the exception of the element *i*:

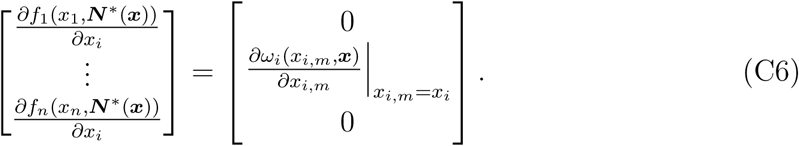

This results in the following equivalence between the evolutionary gradient and the abundance gradient:

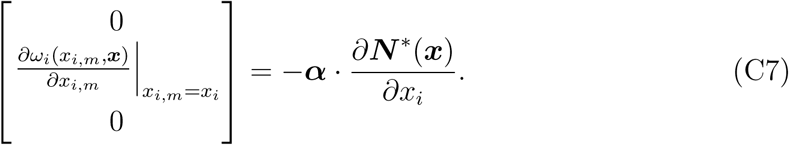

This equation is the generalization to polymorphic (or species) coevolution of the equation (B4) of the monomorphic case. As the matrix *α* represents the linearization of the interaction strength at the equilibrium point, we can assume the matrix not to be singular (i.e., det *α* ≠ 0). This equation shows the equivalence between abundance optimum and singular strategies.

Following the same line as in the monomorphic demonstration, we compute the cross derivative of the relative fitness for each morph *i*,

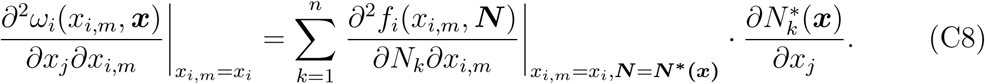

As a singular strategies corresponds to abundance optimum, the cross derivatives equal zeros. This demonstrates that a singular strategy can either be convergent and evolutionary stable (CSS), or non-convergent and evolutionary non-stable (repellor).

Finally, the second derivative relation (equation B9 in the monomorphic case) generalizes to the polymorphic case as

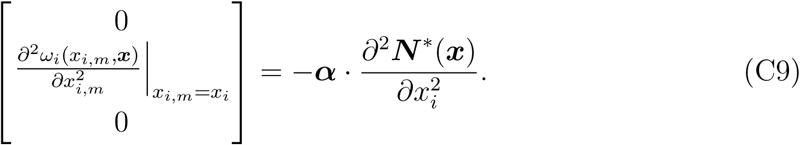

This equation is more complicated as the simple proportional relation of the monomorphic case (equ. B9) and, in general, we cannot guarantee that a CSS reaches a maximum abundance, or that repellors have minimum abundances.

### Class II fitness functions; non-optimization

The main difference with the class I of relative function is that in the vector

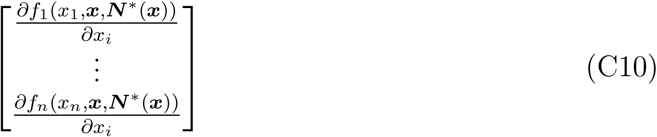

none of the terms

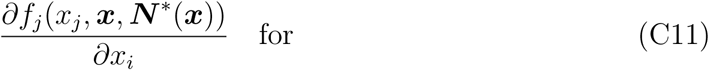

vanish in general. This is so, because of the presence of the term ***x*** in the relative fitness function. That is, in general,

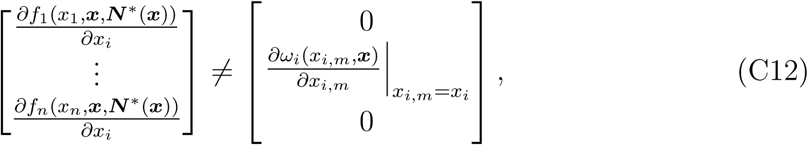

consequently, coevolution does not lead to abundance optimization and any type of singular strategies is possible.

